# Localizing Brain Function Based on Full Multivariate Activity Patterns: The Case of Visual Perception and Emotion Decoding

**DOI:** 10.1101/2021.04.04.438425

**Authors:** Isaac David, Fernando A. Barrios

## Abstract

Multivariate statistics and machine learning methods have become a common tool to extract information represented in the brain. What is less recognized is that, in the process, it has become more difficult to perform data-driven discovery and functional localization. This is because multivariate pattern analysis (MVPA) studies tend to restrict themselves to a subset of the available data, or because sound inference to map model parameters back to brain anatomy is lacking. Here, we present a high-dimensional (including brain-wide) multivariate classification pipeline for the detection and localization of brain functions during tasks. In particular, we probe it at visual and socio-affective states in a task-oriented functional magnetic resonance imaging (fMRI) experiment. Classification models for a group of human participants and existing rigorous cluster inference methods are used to construct group anatomical-statistical parametric maps, which correspond to the most likely neural correlates of each psychological state. This led to the discovery of a multidimensional pattern of macroscale brain activity which reliably encodes for the perception of happiness in the visual cortex, lingual gyri and the posterior perivermian cerebellum. We failed to find similar evidence for sadness and anger. Anatomical consistency of discriminating features across subjects and contrasts despite the high number of dimensions suggests MVPA is a viable tool for a complete functional mapping pipeline, and not just the prediction of psychological states.

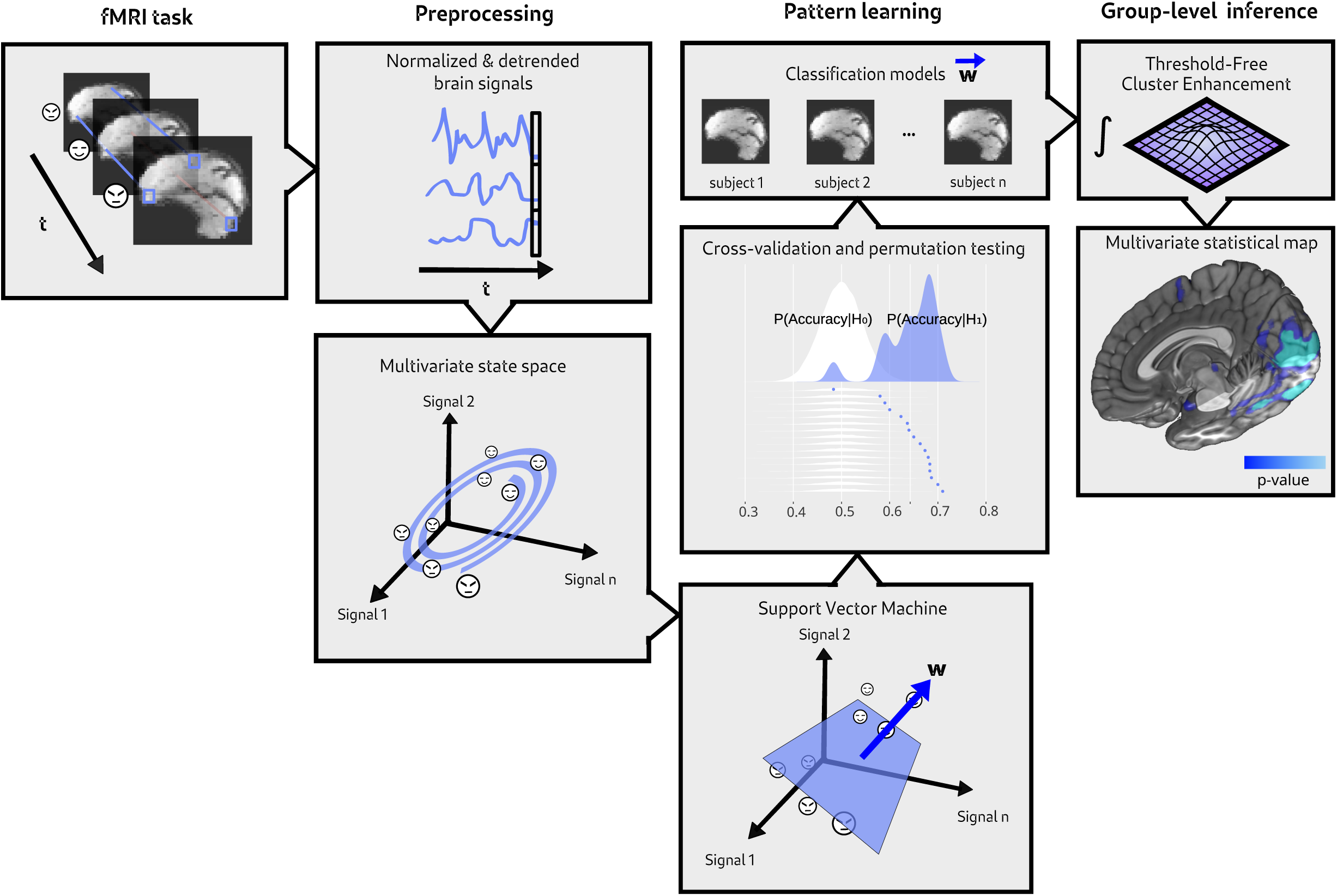

## 1. Introduction

The mapping of segregated brain functions is far from a settled methodology. Take for instance the choice of statistical model in task-oriented functional neuroimaging for modalities like fMRI and PET: while the venerable *mass-univariate analysis* fits separate models to encode each brain time series based on experimental variables, *multivariate pattern analysis* (MVPA) reverses this — particularly multivariate pattern learning — fitting a single model to decode experimental conditions out of the joint activity of several brain signals. The former is excellent at uncovering simple correlations between loci and functions, whereas the latter can potentially provide increased sensitivity due to emergent informational dependencies (Wang et al, 2007; Huettel et al., 2009; Jolly and Chang, 2021), at the expense of computational complexity (De Martino et al., 2008).

In this vein, popular wisdom often dictates that multivariate searches should be limited in one way or another, via dimensionality reduction or restriction to regions of interest (ROI) (Cox and Savoy, 2003; Haynes and Rees, 2005; Kamitani and Tong, 2005) or moving searchlights (Kriegeskorte et al., 2006; Björnsdotter et al., 2011). Furthermore, unlike with the classical mass-univariate approach, the ability to map model parameters onto anatomy may not always be possible, depending on the transparency of the algorithm and its nonlinearities (Schmah et al., 2010).

Paradoxically, there is an enormous interest in studying how the holistic interactions of large-scale brain networks might subserve complex behaviors and mental processes, not just at rest but during tasks (Prinz, 2006; Chialvo, 2010; Avena-Koenigsberger et al, 2018). In fact, most of the recent breakthroughs in statistical learning have been achieved by embracing the high-dimensional nature of problems (Sejnowski, 2020), and safeguards against exaggerated (i.e., over-fitted) findings, such as stringent observation of cross-validation and ROC-curve standards, are now commonplace (Mahmoudi et al., 2012). Moreover, a brain-wide, mappable and statistically sound methodology for functional localization would complete the spectrum of available complementary analytical perspectives purported since the introduction of MVPA (Raizada et al., 2010, Gaonkar and Davatzikos, 2013; Jolly and Chang, 2021). Just as allowing for multivariate dependencies in the first place might provide a whole new picture (Wang et al., 2007; Huettel et al., 2009; Mahmoudi et al., 2012; Lewis-Peacock and Norman, 2013), so could multivariate data drawn from different spatial (or temporal) scales (Hoel et al., 2013; Jolly and Chang, 2021). An auxiliary motivation simply comes from trying to avoid selection bias (Kriegeskorte et al., 2009) and the caveats of dimensionality reduction (Kherif et al., 2002; Wang et al., 2007; Palo et al., 2021).

Yet the merits of such relatively straightforward analysis have seldom — if ever — been fully put to test. Pieces are scattered here and there in the literature: one year after introducing the use of general linear models (GLM) for per-voxel analysis (Friston et al. 1994), Friston et al. (1995) applied multivariate analysis of covariance on whole-brain data, reduced to a space of 35 eigenimages. Ever since, the existing whole-brain experiments either keep constructing a low-dimensional state-space from such multivariate methods like principal components analysis (McIntosh et al., 1996; Mørch et al., 1997; McKeown et al., 1998; Kjems et al., 2002; Kherif et al., 2002; Carlson et al., 2003; LaConte et al., 2003; Wang et al, 2007) or brain atlases and parcellations (Zhang et al., 2020). Another approach is to select individual voxels through (admittedly self-defeating) univariate (Polyn et al. 2005; De Martino et al., 2008; Mwangi et al., 2014) or multivariate means (Craddock et al., 2009). A few works have performed brain-wide multivariate searches without dimensionality reduction, while also failing to translate machine learning models into proper statistical parametric maps (Mourão-Miranda et al., 2005; LaConte et al., 2005; Polyn et al. 2005; Schmah et al., 2010; Raizada et al., 2010).

Raizada et al. (2010) correctly identified and tried to fill the gap, by using GLM in a truly multivariate way. The approach proved promising keeping track of behaviorally-separable linguistic groups, although correction for multiple comparisons was absent in their final statistical-anatomical maps. Gaonkar and Davatzikos (2013) explicitly identified the gap again, and showed how to analytically approximate null models so that statistical significance maps can be derived from model parameters in the case of support vector machine (SVM) classifiers, which they applied to a lie-detection fMRI dataset. To the advantage of the multivariate approach, the False Discovery Rate of statistical maps was shown to be much lower than in the univariate methodology, but this post-hoc discovery was not incorporated to correct brain maps for multiple comparisons.

Here we tested pattern classification analysis as a methodology for anatomical localization in brain-wide, high-dimensional fMRI data. We employed linear SVM classifiers — a relatively interpretable multivariate algorithm, from which a spatial discriminative map can be extracted and directly related to the effect of different experimental conditions on brain activation (Mourão-Miranda et al., 2005; LaConte et al., 2005; Wang et al, 2007; Gaonkar and Davatzikos, 2013). This was tested on a sample of 16 human participants who performed a face perception task. Our goal is to provide anatomical statistical significance maps which are properly corrected for multiple comparisons, and for this we employ a state-of-the-art cluster-informed statistic well-known to the neuroimaging community. Moreover, our task is designed to probe the analysis against variable levels of decoding difficulty: from simple visual stimulation to face detection, to the more ethereal perception of three different emotions. We evaluated whether individual classifiers can learn to predict task state above empirically-estimated chance performance, and whether individual models converge on what the most relevant neural correlates of each cognitive ability are at the group level. We also provide comparisons to the classical mass-univariate (GLM) analysis.

The analysis is expected to reveal well-known early visual cortical areas in contrasts intended to capture the effect of visual stimulation, and components of the so-called face processing network in the ventral stream during face perception (Haxby et al., 2000; Haist and Anzures, 2017). If successful, this would provide greater confidence when pitching the method against a genuinely harder and open problem; namely, segregating and identifying emotions in the brain.

Although affective neuroscience has been fruitful in identifying the anatomical components of the emotional peripheral and central nervous systems; ample disagreement still exists on how to physiologically characterize particular emotional experiences, even among meta-analytical reviews (Vytal and Hamann, 2010; Lindquist et al., 2012; Hamann, 2012; Kragel and LaBar, 2014; Guillory and Bujarski, 2014; Kragel and LaBar, 2016; Celeghin et al., 2017). It is unclear how the distributed activity of many limbic and other mid-line structures, from the posterior perivermian cerebellum to the medial prefrontal cortex (among others), gives rise to phenomena such as sadness, rage or positive hedonic valence. This realization has in turn brought multivariate methods into emotion research (Rolls et al., 2009; Baucom et al., 2012; Chikazoe et al., 2014; Shinkareva et al., 2014; Chang et al., 2015; Sitaram et al., 2011; Kassam et al., 2013; Saarimäki et al., 2015; Kragel and LaBar, 2015) and emotional perception research. In general, by use of ROIs and searchlights, localized multivariate activity has been found to outperform its univariate counterpart distinguishing among emotions (see Table 1). However, this has not been extended to a proper brain-wide search, insofar as face perception is concerned.

**Table 1:** Survey of experimental studies on emotion perception that employed MVPA. Column descriptions: Modality: stimuli modality. Emotions: emotions under investigation (usually supplemented with an extra “neutral” category). ROI: region of interest. Accuracy: average classification accuracy, compared to theoretical random accuracy given the number of emotion categories. SVM: support vector machine. SMLR: sparse multinomial logistic regression. MGPC: mixture of Gaussian processes for classification. RSA: representation similarity analysis.

| *Reference* | *Modality* | *Emotions* | *ROI* | *Algorithm* | *Accuracy* |
| --- | --- | --- | --- | --- | --- |
| *Pessoa and Padmala (2007)* | *visual* | *fear* | *many* | *SVM* | *78% > 50%* |
| *Ethofer et al. (2009)* | *auditory* | *happiness, anger, sadness, relief* | *auditory cortex* | *SVM* | *33% > 20%* |
| *Said et al (2010)* | *visual* | *happiness, anger, sadness, disgust, surprise, fear* | *superior temporal sulcus, frontal operculum* | *SMLR* | *22% > 14%* |
| *Peelen et al. (2010)* | *auditory, visual (faces and body language)* | *happiness, anger, sadness, disgust, fear* | *brain (searchlight)* | *RSA* | *N.A.* |
| *Kotz et al (2012)* | *auditory* | *happiness, anger, sadness, surprise* | *brain (searchlight)* | *SVM* | *28% > 20%* |
| *Skerry and Saxe (2012)* | *reading, visual (faces)* | *21 appraisals* | *theory of mind network, brain (searchlight)* | *SVM* | *~8% > 5% (ROIs)* |
| *Wegrzyn et al. (2015)* | *visual (faces)* | *happiness, anger, fear* | *many* | *MGPC* | *~32% > 25%* |

**Table 2:** Demographic features of the 16 successfully included participants. S.D: standard deviation.

| *n=16* |  |  |
| --- | --- | --- |
| *Age (years)* | $\bar{x}$ | *S.D.* |
|  | 25 | 3.01 |
| *Sex* | *Female* | *Male* |
|  | 8 | 8 |
| *Education level (obtained or in progress)* | *Undergraduate* | *Postgraduate* |
|  | 8 | 8 |

## 2. Materials and Methods

### 2.1 Sample

Sixteen volunteers from both sexes (8 female, 8 male) were recruited at UNAM campus Juriquilla from October 2019 to June 2020. Participants were briefly interviewed to exclude those previously diagnosed with neurological or psychiatric conditions. All of them reported having right-handed phenotype with the exception of one male. Prior to the study, subjects formally consented to participating after being informed of its aims, risks and procedures — in accordance with the 1964 Declaration of Helsinki — and were compensated with their brain scans and free diagnostics by a radiologist.

### 2.2 Image acquisition

Images were obtained from a 3-Tesla General Electric Discovery MR750 scanner at the Magnetic Resonance Unit at UNAM campus Juriquilla, during a single session per participant. The protocol included five echo-planar imaging (EPI) blood-oxygenation-level-dependent (BOLD) sequences for fMRI, 185 volumes each. A T1-weighted (T1w) scan of head anatomy was also acquired. Sequence parameters are described in Table 3. Electromagnetic responses were recorded using a head-mounted 32-channel coil.

**Table 3:** Sequence parameters used for the MRI protocol. Abbreviations: T1w: T1-weighted, FSPGR: fast spoiled-gradient echo, TInv: inversion time parameter for FSPGR T1w imaging.

| *Parameter* | *EPI BOLD* | *T1w FSPGR* |
| --- | --- | --- |
| *Slice orientation* | *Axial* | *Axial or sagittal* |
| *Slices* | *35* | *176* |
| *Field of view* | *64×64* | *256×256* |
| *Voxel size* | *(4 mm)<sup>3</sup>* | *(1 mm)<sup>3</sup>* |
| *Flip angle* | *$\pi/2$* | *$3\pi/45$* |
| *TR (ms)* | *2000* | *8.18* |
| *TE (ms)* | *30* | *3.19* |
| *TInv (ms)* | *N.A.* | *450* |

### 2.3 Stimuli and task

Each of the five fMRI sequences was temporally coupled to a psychological block-based task implemented in PsychoPy 3.0.1 (Peirce, 2007). All five tasks were identical, save for the pseudo-random order in which their 30 s blocks were administered. A total of six block classes were used: happy faces, sad faces, angry faces, neutral faces, pseudo (scrambled) faces and low-stimulation. Neutral/inexpressive faces might provide an extra control when contrasting among emotions. Pseudo-faces and dim blocks were introduced so as to buttress and diagnose the analysis pipeline, by way of more trivial contrasts (like pseudo-faces vs low-stimulation and faces vs pseudo-faces).

Each block, in turn, comprises 10 randomly-presented images belonging to that class, each one shown for about 3 s and without possibility of re-instantiation during the same block. Each block occurs twice per sequence, yielding a total of 12 of them (360 s = 6 min). After their presentation, participants had to wait for 10 s before concluding the sequence, in order to capture the hemodynamic response (HR) elicited by the last stimuli. A selection of 10 gray-scale photographs per category of frontal human faces (male and female) served as stimuli. These were chosen from the “Pictures of Facial Affect” database (Ekman, 1976). As for the low-stimulation (a.k.a. “dim”) blocks, a small but visible fixation cross was made to fluctuate from quadrant to quadrant at random every 3 s. The whole task is summarized in Figure 2.

**Figure 1:**
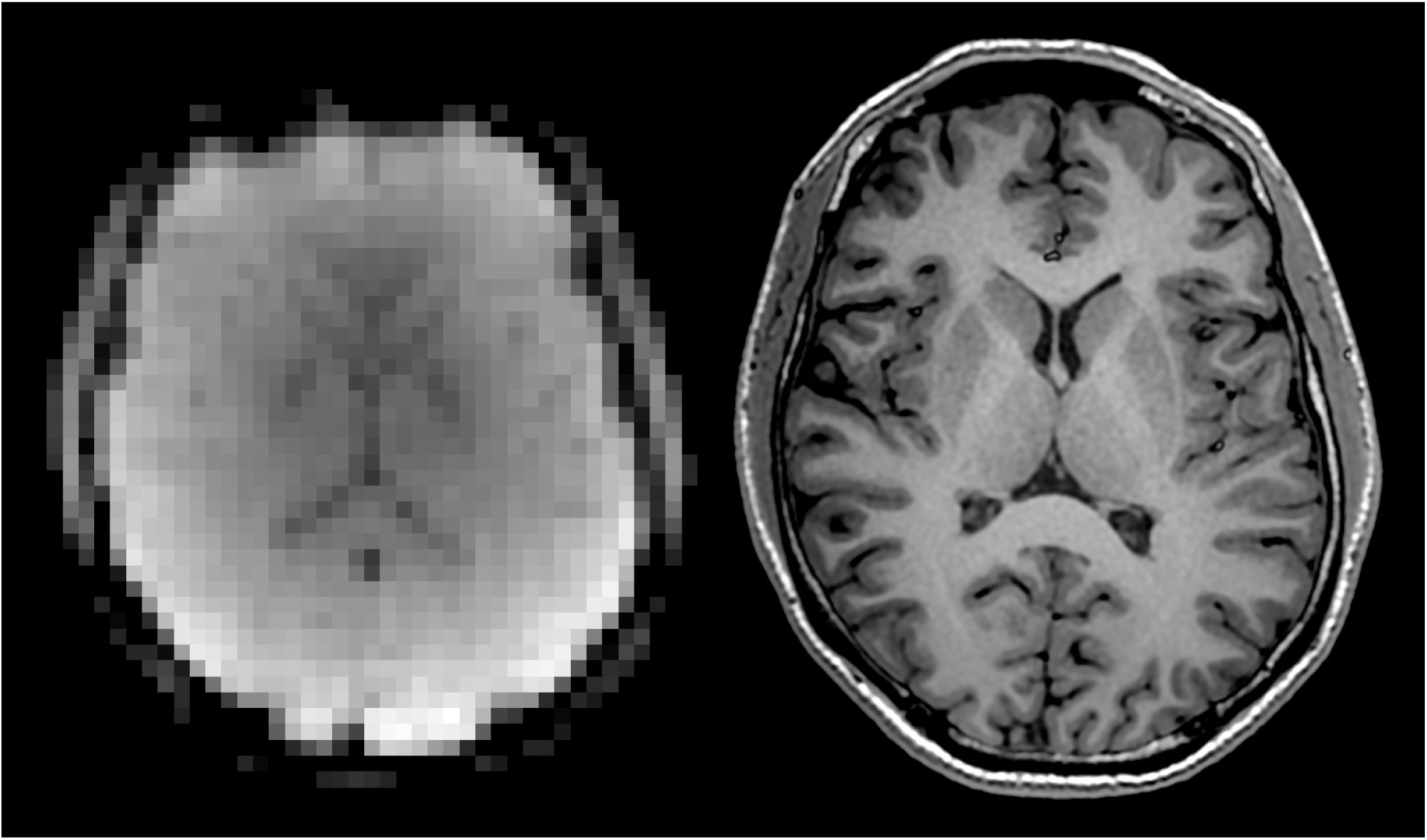
Raw samples of both image modalities for a single subject in our dataset (in the same order as the columns in *Table 3*).

**Figure 2:**
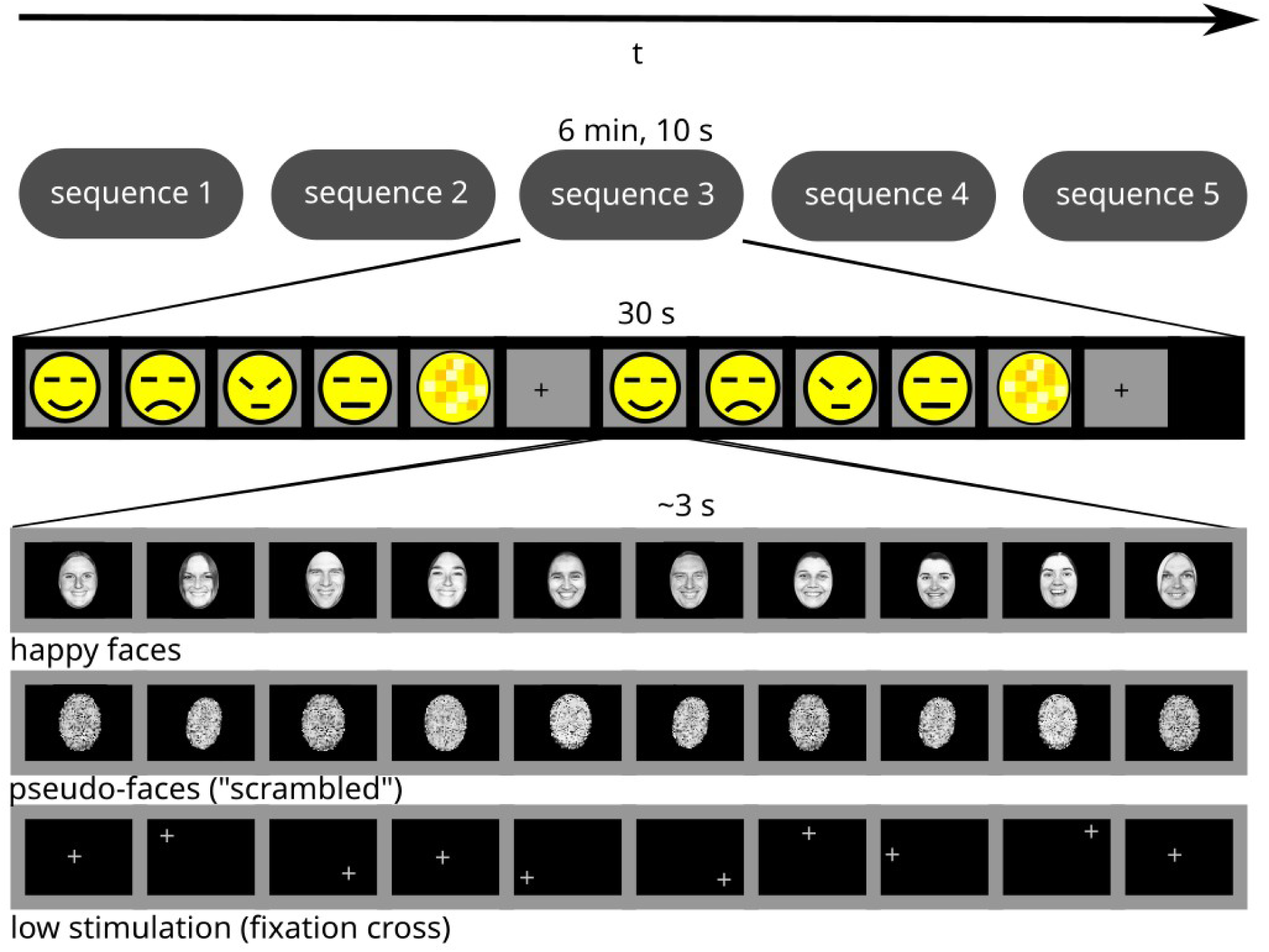
Block-paradigm design of the psychological experiment. The horizontal axis corresponds to the passing of time. Rectangles represent stimulation units: sequences (top row), blocks (second row) or individual stimuli.

Additionally, behavioral responses were recorded throughout the task in order to measure performance and thus evaluate the suitability of physiological data for further analysis. Participants were given an orthogonal task: to indicate whether faces belonged to a man or a woman as soon as they were perceived. The response was submitted with the press of a button, one in each hand. Analogously, for scrambled and dim blocks (when no faces should have been perceived), the instruction was to simply report image change, alternating between buttons. In this fashion, motor activity remained rather homogeneous for all blocks, minimizing a possible confounding effect when contrasting among faces and pseudo-faces. Statistical analysis of behavioral data was conducted using the R programming language.

### 2.4 Analysis methods

#### 2.4.1 Code and data accessibility

All the software necessary for reproducing, analyzing or adapting the present study is made available as free software and may be downloaded at https://github.com/isacdaavid/np-mvpa . Original data and final group activation maps in standard space may be downloaded respectively from OpenNeuro (https://doi.org/10.18112/openneuro.ds003548.v1.0.1) and Neurovault (https://identifiers.org/neurovault.collection:9492).

#### 2.4.2 Image preprocessing

T1w images were submitted to the volBrain tissue segmentation and volumetry web-based service (Manjón and Coupé, 2016). The resulting brain and gray/white matter masks we used for deskulling the field-bias-corrected T1w images and, later on, for selection of fMRI voxels. fMRI sequences were concatenated by temporal order in one long sequence per subject, then the result underwent the following preprocessing pipeline due to the FSL 6.0 utilities (Jenkinson et al., 2012): high-pass frequency filter (>50 s) and interpolation for slice-time correction (interleaved acquisition) (Woolrich et al., 2001), affine movement correction and co-registration (Jenkinson and Smith, 2001; Jenkinson et al, 2002) with the respective T1w anatomical reference and the standard MNI-152 T1w template (Fonov et al., 2009; Fonov et al., 2011) at 1 mm of resolution. After registration, the corresponding resulting matrices were applied to the volBrain masks, so as to transform them to the low-resolution subject-space of fMRI images. Gray matter time series were extracted afterwards (about 10000 time series, depending on the subject), and linear trends were subtracted by preserving residuals from a simple linear regression performed on each of the five sequences that comprise the long concatenated time series. Finally, and seeking not to bias classification models in any dimension, the composite time series at each voxel was normalized to z-scores.

#### 2.4.3 Univariate analysis

To assess the feasibility and performance of the brain-wide multivariate approach against the golden standard in functional brain mapping, we investigated the same two-way contrasts included in multivariate analysis using FSL 6.0 in a classical mass-univariate analysis. These contrasts are grouped into *visual stimulation* (“dim vs scrambled”, “dim vs neutral”, “dim vs angry”, “dim vs sad”, “dim vs happy”), *face perception* (“scrambled vs neutral”, “scrambled vs happy”, “scrambled vs sad”, “scrambled vs angry”) and *emotion perception* (“angry vs happy”, “sad vs happy”, “sad vs angry”, “angry vs neutral”, “happy vs neutral”, “sad vs neutral”). Preprocessed data (up until subtracting linear trends and normalizing) were spatially-smoothed with a Gaussian convolution kernel of 5 mm FWHM. Each of the six block classes described for the task was considered as a column-vector regressor in the design matrix ***X***, after convolving them with a zero-lag, double-gamma HR curve. ***X*** was augmented with the time derivatives of each convolved regressor, but no motion covariates were added. GLMs were fitted afterwards. GLM is a matrix-form extension to multiple linear regression, which models each physiological time series (column ***Y***) as a linear combination of ***X*** plus some Gaussian error ***E***. The model reads (Mahmoudi et al., 2012):

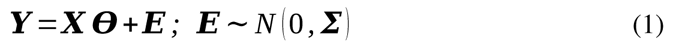

Assuming trial independence and homocedasticity, maximum likelihood estimation or ordinary least-squares estimation may be followed to obtain the so-called normal equation, which optimizes parameters ***θ*** according to:

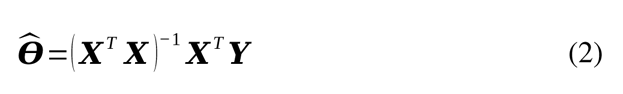

After estimation, contrasts of parameter estimates are subjected to the same procedure as the multivariate model parameters for purposes of group-level inference. This is described in more detail in the *2.4.5 Group inference* subsection.

#### 2.4.4 Multivariate pattern analysis

Once preprocessed as described in subsection 2.4.2, multivariate decoding of fMRI patterns was conducted using the pyMVPA 2.5.0 Python library (Hanke et al., 2009). We trained a linear SVM classifier per subject and block contrast combination using all available brain volumes (120 volumes per class for the training phase, 30 for testing). In addition to the contrasts described in the previous section, emotion-related ones were augmented with the _(_^4^_3)_ possible three-way classification problems and the single four-way contrast. The supervised SVM algorithm learns a hyperplane for binary classification in high-dimensional state space (Vapnik and Chervonenkis, 1974; Boser et al., 1992). Given a vector ***w*** orthogonal to the hyperplane, the SVM decision rule is equivalent to the sign of the projection of unseen data vectors ***Y*** *_i_* on ***w***, adding or subtracting the necessary constant *b* so as to make the result exactly 0 at the hyperplane:

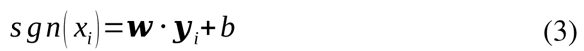

Out of all possible hyperplanes, SVM’s key insight is to estimate the one that maximizes separation margin to the most difficult training data: the support vectors right above opposite margin lines. Since margin width can be calculated from pairs of positive-class and negative-class support vectors according to:

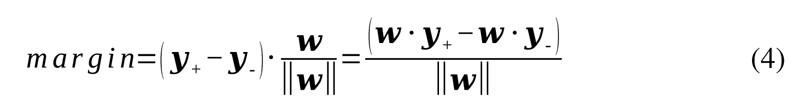

by constraining the decision rule to satisfy _|_***w*** *⋅* ***Y*** *_i_*+*b*_|_*≥* 1 or similar criteria and substituting on equation (4), one can show that maximizing the margin — and therefore obtaining an optimal model — is equivalent to a quadratic programming problem with ‖***w***‖ as the cost function to be minimized (mathematical details are discussed by Mahmoudi et al (2012)):

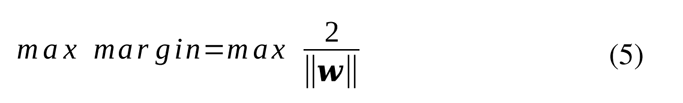

SVM models were cross-validated using each of the five sequences as a fold, and their mean classification accuracy (hits / misses) served as a summary test statistic. This was compared against an empirical null model in a non-parametric rank-based test, by estimating the probability distribution of average classification accuracy given *H_0_* via Monte-Carlo simulations: surrogate data is computed by randomly shuffling the class labels of the training data partition 5000 times, and five-fold cross-validation is again conducted for each permutation. Then, a p-value for that particular subject and class combination can be calculated as the proportion of random results equal to or greater than the original classification accuracy.

We also explored the effect of different delays between stimulus onset and volume labeling (from 0 s to 10 s, every 2 s), instead of assuming a single optimal HR peak (Lewis-Peacock and Norman, 2013); although, based on common practice and prior knowledge about typical HR, a labeling delay of 4 s was fixed a priori when conducting all group-level statistical inferences.

#### 2.4.5 Group inference

To assess whether individual models reveal generalizable anatomical structures at the group level we employed yet another non-parametric test, one per contrast. In particular, we used FSL 4.0’s *randomise* (Winkler et al., 2014) with 5000 rounds of the Threshold-Free Cluster Enhancement (TFCE) (Smith and Nichols, 2009) operating on the group of SVM weight vectors to generate null data and then perform a two-tailed test (models were spatially smoothed in advance with a 5 mm FWHM Gaussian kernel, and to make them comparable, they were transformed to the standard 1 mm MNI-152 space, as well as normalized so that all weight vectors become unitary). In the case of the univariate models, the only difference is that TFCE receives a group of GLM contrasts of parameters as input data (i.e., also in a two-tailed test with both positive and negative effects, prior spatial smoothing and transformation to the 1 mm MNI-152 space).

Given an anatomical image *h(v)* with scalar values representing model parameters (or contrasts among them), the TFCE statistic at some voxel *v* is defined as the integral (in the Lebesgue sense) of cluster size *s(v, h)* times the cluster-defining “height” *h*:

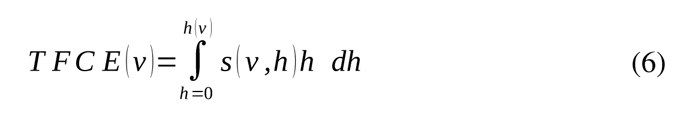

In practice, equation (6) is modified for fMRI and EEG data, where the default is to favor *h*, squaring it, while taking the square root of *s* (Smith and Nichols, 2009). Since all possible cluster-forming thresholds are considered by the integral, TFCE is generally regarded as a more principled alternative to other nonparametric cluster-informed inference methods, while still providing strong family-wise error (FWE) control (Roiser et al., 2016), as expected of nonparametric approaches. For the sake of visualization, we set a cluster cutoff value of 0.01 in the corrected p-value brain maps to render voxels that are unlikely to correlate with the task by pure chance. Brain structures were labeled by agreement of the AICHA (Joliot et al., 2013), AAL (Tzourio-Mazoyer et al., 2022) and Harvard-Oxford (Desikan et al., 2006) atlases. The same unthresholded maps may be downloaded from https://identifiers.org/neurovault.collection:9492 .

The explicit validation of multivariate models produces an extra piece of information, in the form of performance statistics (e.g., classification accuracy). We report individual and group-average p-values (one per contrast) on classification accuracy, as obtained in per-subject permutation tests, as well as estimated group effect size (one per contrast) by comparing classification accuracies to the null distribution according to Cohen’s *D* statistic. Echoing recommendations from the statistical community, we did not set arbitrary “significance” thresholds, but rather regard these values as lying on a continuous evidence scale (Wasserstein and Lazar, 2016; Amrhein and Greenland, 2018; Wasserstein et al, 2019).

## 3. Results

### 3.1 Behavior

See the attached supplementary material.

### 3.2 Visual stimulation and face perception

All contrasts meant to distinguish between high and low visual stimulation and between face and pseudo-face perception presented strong evidence of successful decoding using the multivariate model, both on an individual and on a group-level basis. In the case of visual stimulation, all p-values on classification accuracy per contrast (both individual and average) were found to be lesser than _2_ *_⋅_* _10_*^−^* ^4^: the smallest result that could have been obtained with 5000 permutations. Moreover, we found extremely large group effects; with Cohen’s D ranging from *D*=6.5 (dim vs neutral) to *D*=7.6 (dim vs happy). Similarly, face perception compared to its baseline always resulted in a *_p_*_<2_ *_⋅_* _10_*^−^* ^4^; both individually and as group means. Cohen’s statistic ranged from *D*=5.2 (scrambled vs neutral) to *D*=6.8 (scrambled vs angry). Figure 3 displays hypothesis tests for a couple of experimental contrasts, for the sake of illustrating the nature of results. Statistical maps of model parameters are presented in Figure 4.

**Figure 3:**
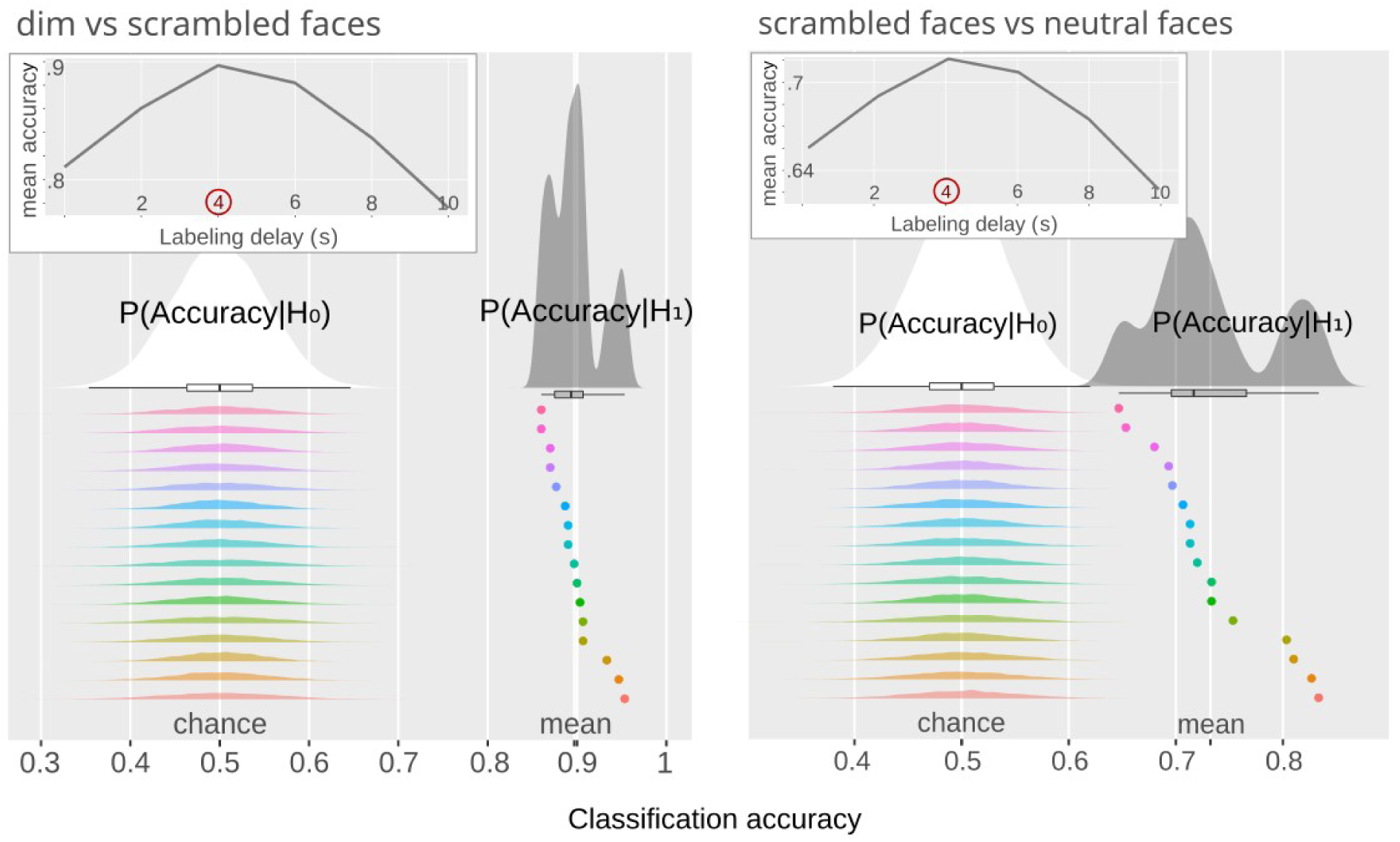
All nine contrasts related to simple visual perception and face perception (only two shown here) are strongly dissociable, according to their brain-wide activity patterns. Embedded at the top-left corner of each subfigure: group mean classification accuracy as a function of labeling delay. Greater subfigure: hypothesis tests of classification accuracy for a preset labeling delay of 4 s after stimulus onset. Rainbow-colored dots at the bottom stand for the cross-validated classification accuracy of each participant, compared to their respective estimated null distributions.

**Figure 4:**
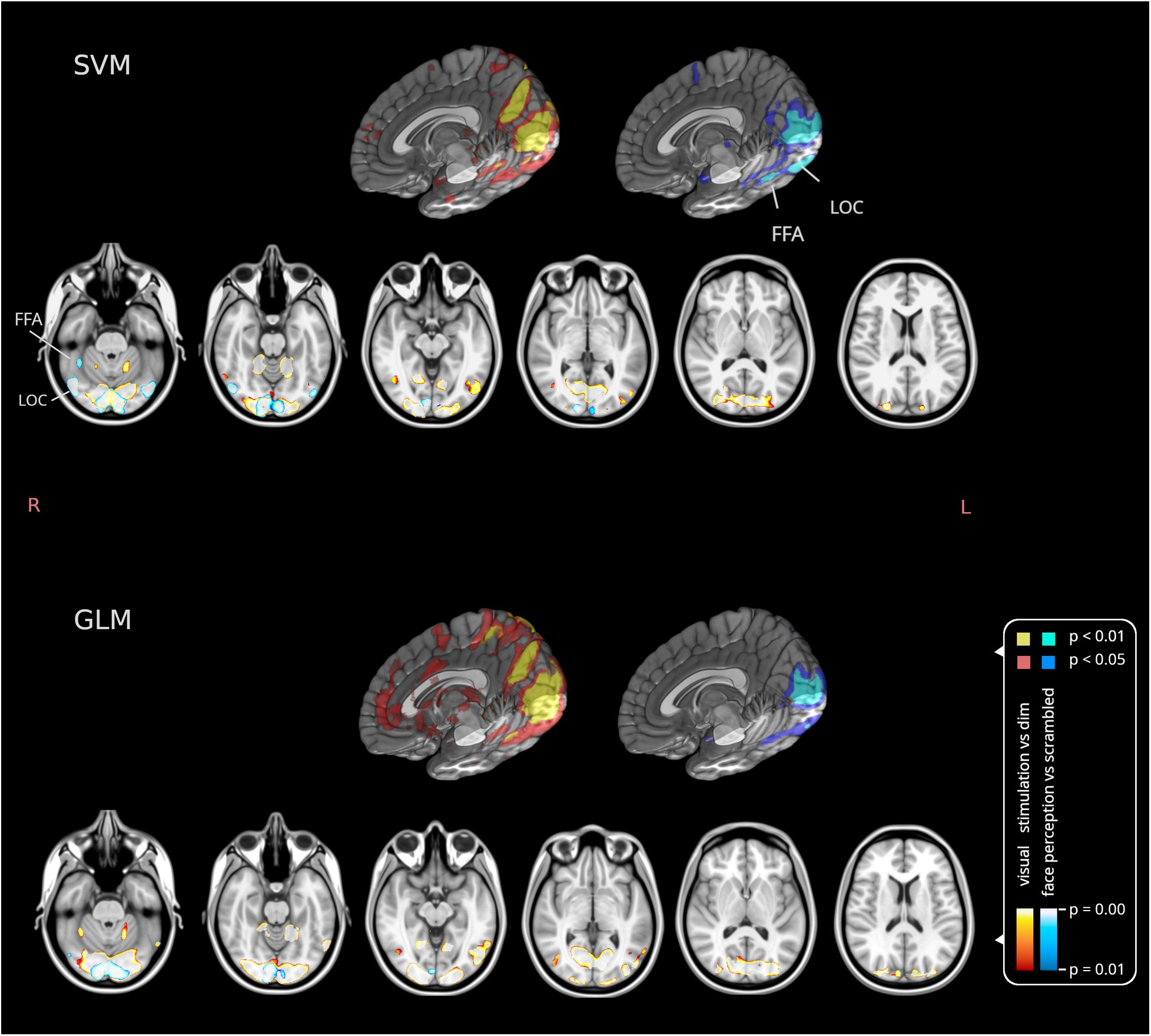
Anatomical distribution of thresholded (p_FWE_ < 0.01) statistical maps according to the group-level TFCE analysis. Yellow: visual stimulation (average of FWE-corrected group p-values over related contrasts). Cyan: the same average of FWE-corrected group-level p-values, but for face perception-related contrasts. Two views of the same data are offered: 3D rendering of right hemisphere (on top), axial slices in radiological orientation (bottom). In both cases results are imposed on the T1w MNI-152 template. Unthresholded brain maps may be downloaded from https://identifiers.org/neurovault.collection:9492.

Both univariate and multivariate approaches agree on two prominent, bilaterally-symmetric occipital clusters whose activity correlates with the presentation of visual stimuli (see Figure 4 and Table 4): one posteromedial, encompassing part of the primary visual cortex (V1) and extra-striate areas like V2, possibly reaching parts of ventral V3 (VP) and color and form-related V4. From there it crosses parenchymal boundaries to the medial posterior cerebellar lobe, in its anterior portion. The second major cluster lies at the anterior medial occipital cortex, and in the univariate model this is due to anticorrelation. It includes a calcarine component at anterior V1, part of the V2 ring and then extends more dorsally into the cuneus to motion-related V3a and the midline section of form-related V3. Slightly above the cutoff p-value threshold, associated activity also survives bilaterally into dorsal stream regions (precuneus Brodmann area 7 and V5/MT) according to both univariate and multivariate models, perhaps because of an unintended effect of having a moving fixation cross.

**Table 4:** Summary of clusters shown for *Figures 4 and 7*, according to the group level analysis of multivariate models, together with representative minima and their coordinates. All coordinates follow the Montreal Neurological Institute system. Structures were localized by consensus of the AICHA, AAL and Harvard-Oxford atlases.

| Structures spanned by cluster | Left Hemisphere |  |  |  |  | Right Hemisphere |  |  |  |  |
| --- | --- | --- | --- | --- | --- | --- | --- | --- | --- | --- |
|  | Cluster size | p<sub>FWE</sub>-value | x | y | z | Cluster size | p<sub>FWE</sub>-value | x | y | z |
| Visual stimulation |  |  |  |  |  |  |  |  |  |  |
| Posterior Calcarine Sulcus (V1, V2), Perivermian Posterior Cerebellar Lobe (PPCL) | 20812 | 0.0002 (V1, V2) | -11 | -92 | 0 | 23442 | 0.0002 (V1, V2) | 11 | -92 | 0 |
| Anterior V1, V2, V3a, V3, Perivermian Anterior Cerebellar Lobe (PACL) | 19682 | 0.0002 (PACL) | -16 | -52 | -22 | 24024 | 0.0002 (PACL) | 19 | -53 | -21 |
|  |  | 0.0002 (V1, V2) | -10 | -71 | 13 |  | 0.0002 (V1, V2) | 10 | -71 | 13 |
| Face perception |  |  |  |  |  |  |  |  |  |  |
| Posterior calcarine sulcus (V1, V2) | 11425 | 0.0002 (V1,V2) | -7 | -92 | -10 | 12545 | 0.0002 (V1,V2) | 7 | -92 | -10 |
| Middle Occipital Gyrus (MOG), Posterior and Middle Fusiform Gyrus (FFA) | 4855 | 0.0002 (MOG) | -46 | -74 | -12 | 8204 | 0.0002 (MOG) | 46 | -74 | -12 |
|  |  |  |  |  |  |  | 0.0002 (FFA) | 39 | -49 | -26 |
| Emotion perception (happy vs neutral) |  |  |  |  |  |  |  |  |  |  |
| Perivermian Posterior Cerebellar Lobe (PPCL), Posterior Calcarine Sulcus (V1, V2) | 18768 | 0.0002 (PPCL) | -23 | -76 | -32 | 19367 | 0.0002 (PPCL) | 29 | -73 | -27 |
|  |  | 0.0002 (V1, V2) | -5 | -94 | 0 |  | 0.0002 (V1, V2) | 8 | -90 | 0 |
| Lingual gyrus (LG) | 12363 | 0.0002 (LG) | -18 | -51 | -11 | 24385 | 0.0004 (LG) | 18 | -47 | -10 |

Face perception was related to two quasi-bilaterally-symmetrical clusters: an anticorrelated pattern at V1 and V2. The second one comprises the middle occipital gyrus (MOG) in the lateral occipital cortex (LOC). Thirdly, multivariate analysis consistently revealed the fusiform face area (FFA) at the right inferior temporal cortex as well, with important subthreshold evidence at its left counterpart. In comparison, univariate analysis barely detected a small cluster at LOC; whereas only 2/4 contrasts showed suprathreshold evidence at FFA (also see Figure 4 and Table 4).

### 3.3 Emotion perception

Figure 5 is a compilation of group tests for the remaining 11 emotion-related contrasts in the form of a Venn diagram. We observed wide variation in model success, but structured according to the emotions under probe; from *p*=0.05 and *D*=3.3 (happy vs neutral) to *p*=0.85 and *D*=*−* 1.3 (anger vs sadness). It is suggested from Figure 5 that contrasts which included *happiness* in general outperformed their respective null models (happy vs neutral: p̄=0.05, *D*=3.3; happy vs sad vs neutral: p̄=0.06, *D*=2.4; happy vs angry vs neutral: p̄=0.11, *D*=2.2; happy vs sad: p̄=0.15, *D*=1.8; happy vs angry: p̄=0.16, *D*=2.0; happy vs sad vs angry vs neutral: p̄=0.18, *D*=1.6; happy vs sad vs angry: p̄=0.31, *D*=0.8). On the other hand, classification models that excluded this emotion did not (sad vs neutral: *p*=0.38, *D*=0.4; angry vs neutral: p̄=0.53, *D*=*−* 0.1; sad vs angry vs neutral: p̄=0.56, *D*=*−* 0.4; sad vs angry: p̄=0.85, *D*=*−* 1.3). The importance of happiness driving accurate prediction was further confirmed via qualitative inspection of confusion matrices. Interestingly, performance degraded with increasing number of categories (as expected), unless the classification problem was augmented with the neutral category, in which case performance improved. As an extension of Figure 5, Figure 6 displays the rank-size distributions of individual p-values in more detail.

**Figure 5:**
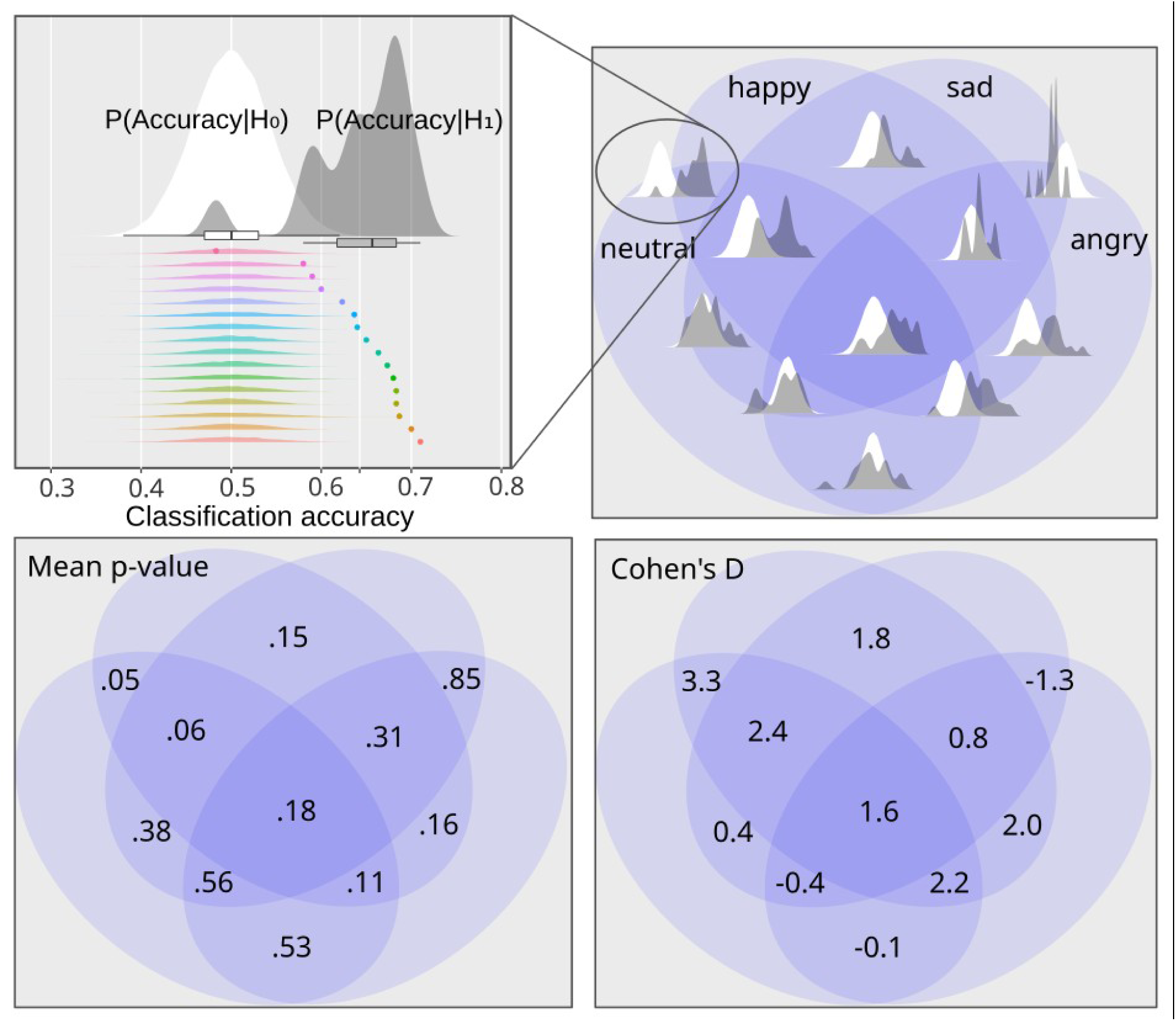
Group aggregates of hypothesis tests on classification accuracy for all emotion combinations, together with their associated mean p-value and effect size (Cohen’s D). Gray and white probability distributions are shown for every intersection in the Venn diagram at the upper right corner. As in *Figure 3*, the gray one is the aggregate of cross-validated classification accuracies when individual models are trained to discriminate the states codified by that part of the Venn diagram; whereas the white probability distribution is the aggregate of corresponding classification accuracies by pure chance, according to a null model (one per subject) estimated with 5000 random permutations of data labels. For the sake of illustration, this is expanded at the upper left corner for a single combination of states (happy vs neutral), with the 16 individual models shown below in rainbow colors (point cross-validated classification accuracy and null distribution composed of 5000 data points).

**Figure 6:**
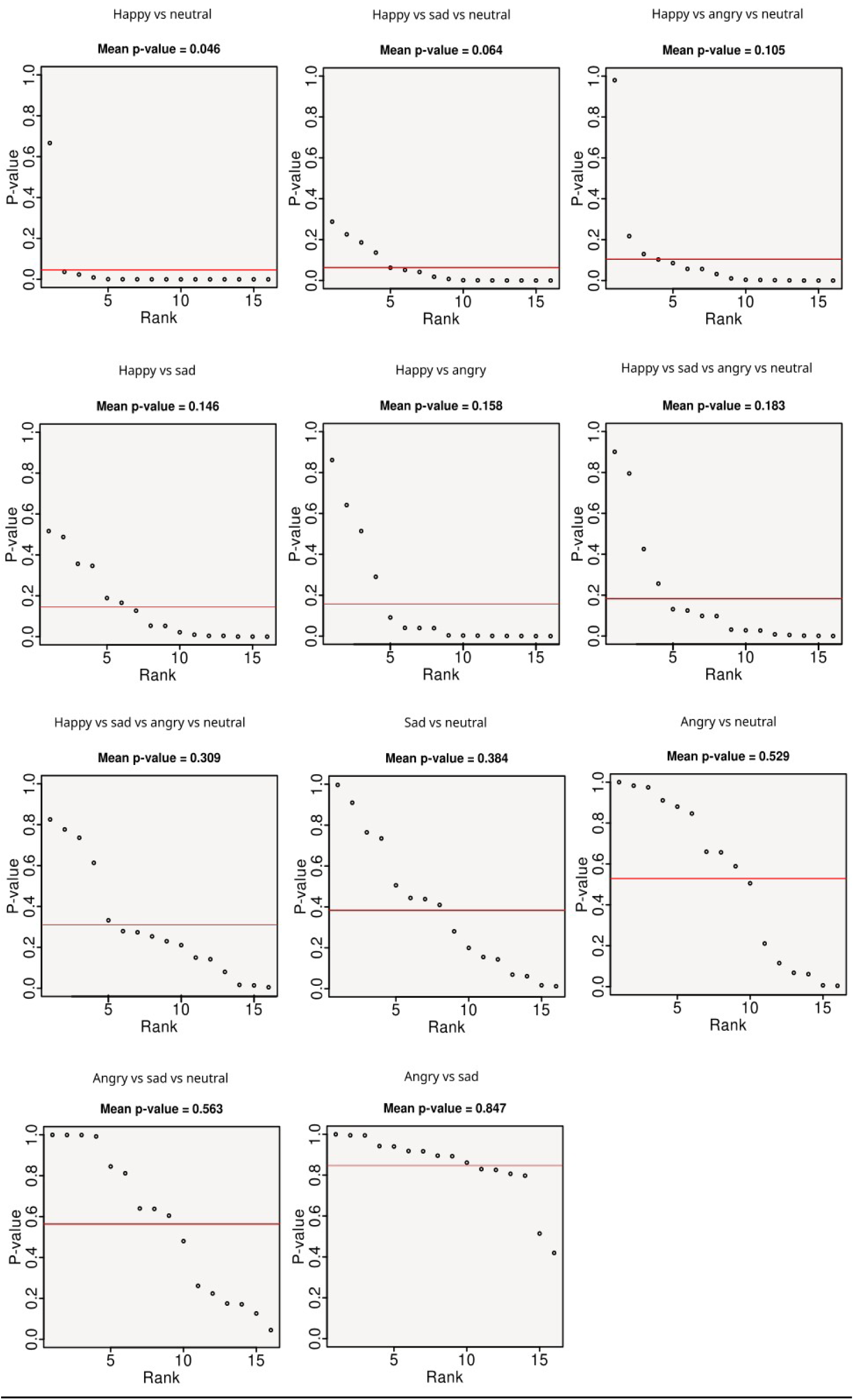
Empirically estimated individual p-values on classification accuracy for each of the 11 emotion contrasts and 16 subjects, ordered by rank, from which mean p-values per contrast are computed for *Figure 5*. The mean p-value is also indicated with a red horizontal line. In general, classification performance improves with the “happiness” and “neutral” face classes.

Group-level inference of SVM parameters resulted in suprathreshold clusters with high co-localization — not just among subjects within the same contrast, but even among contrasts — with two possible characteristic anatomical spans, depending on whether neutral faces had been included in the classification problem. In their absence, detection of the happy faces consistently depended on activity at the occipital pole and its midline and ventral surroundings (posterior V1, posterior V2, ventral V3, V4/ V8) as well as anterior V1 and V2 (both dorsal and ventral). This is shown in yellow clusters in Figure 7. When also faced with the neutral control stimuli (cyan clusters in Figure 7), SVM was forced to extend the search to lower-order and higher-order structures: the pulvinar thalamic bodies, the anterior lingual gyrus (LG) — close to parahippocampal tissue — and the perivermian posterior cerebellar lobe (PPCL). See table 4 for a summary of clusters and minima, together with their stereotactic coordinates. Although not shown, the most prominent subthreshold evidence (*p_FWE_* <.05) was found at the left amygdala and LOC. Subject-level SVM models also gave prominent weighting to the orbitofrontal/ventromedial prefrontal cortex, however, the amount of clusters and their parameter signs in that region were too heterogeneous for evidence to accumulate at the group-level.

**Figure 7:**
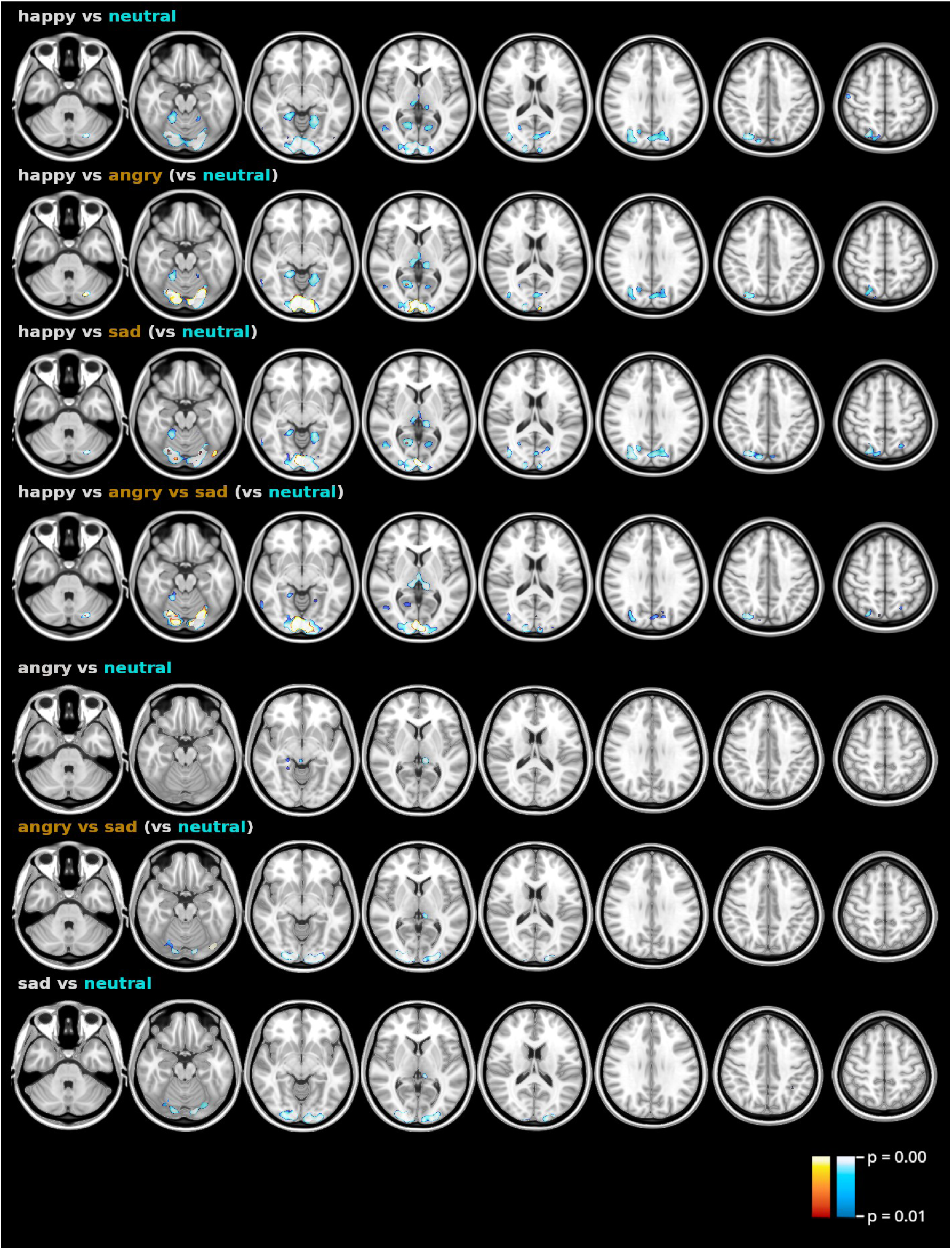
Group anatomical distribution of thresholded (p_FWE_ < 0.01) statistical maps for emotion-related contrasts, as obtained by the TFCE group-level analysis operating on individual SVM parameter vectors. Each row represents one of the 11 possible contrasts, already described in terms of classification accuracy in *Figures 5 and 6*. Contrasts in cyan inclu_2_d_4_e neutral faces as an extra control category in the classification problem. See *table 4* for a description of clusters and minima with coordinates. Slices are in radiological axial orientation in the standard MNI-152 space. Unthresholded maps may be downloaded from https://identifiers.org/neurovault.collection:9492.

By contrast, discrimination of sadness and/or anger (whose decodability was considerably worse, as shown in Figures 5 and 6) resulted in virtually nonexistent anatomical clusters: the left pulvinar body in the thalamus (angry vs neutral) and posterior V1/V2 and PPCL (sad vs neutral). These are rendered in the three bottom rows of Figure 7. Remarkably, no suprathreshold cluster or voxel, correlated or anticorrelated, was found for any of the emotion-related contrasts using classical mass-univariate analysis.

## 4. Discussion

Results validate the feasibility of looking for multivariate correlations between functional neuroimaging and perceptual phenomena of varying complexity, and of turning learned data patterns into statistical-anatomical maps for localization of relevant brain structures. This was shown for an extrinsically high-dimensional state space using all the available gray-matter information, which differs substantially from the routes taken by both univariate analysis and other multivariate studies. It is remarkable that an algorithm as modest as the kernel-less SVM can characterize many psychological states of scientific interest in a purely data-driven approach, to the point of surpassing the classical methodology for its brain mapping features.

Results for the simple visual stimulation subexperiment are almost identical according to both approaches (Figure 4). However, remaining contrasts decidedly favored the brain-wide multivariate analysis. Univariate analysis notoriously failed to consistently discover the right FFA for face perception contrasts, and in terms of the emotions carried by faces, it could not find a single correlated voxel. This suggests that the increased sensitivity of multivariate algorithms is also preserved at the brain-wide level, even in the absence of dimensionality reduction.

That said, it might still seem tempting to disparage results for the emotional subtask, due to major reliance on visual, rather than emotion-related areas other than the perivermian posterior cerebellar cortex (Schmahmann and Sherman, 1997) and the lingual gyrus (which has been involved both in complex visual processing and in emotional and face processing: Isenberg et al, 1999; Kehoe et al, 2013); although as mentioned in the Results section, SVM also relied on the dynamics of the amygdala and the ventromedial prefrontal cortex to drive a decision, albeit less prominently. However, the fact that at least one emotion class (happiness) reliably elicits a distributed activity fingerprint — which was invisible to GLM in the first place — speaks of the advantages of moving beyond simple mass-univariate modeling when it comes to not only neural decoding but also brain mapping. Whether this particular “happy interlocutor state” is truly a non-collateral biological feature of social significance is hard to answer with our data. On one hand, the connectivity and modulation of core affect regions upon the visual cortex has been well-attested by independent studies (Amaral and Price, 1984; Damaraju et al., 2009) and meta-analytic reviews of emotion (Vytal and Hamann, 2010; Lindquist et al., 2012). Moreover, recent experiments using electrophysiological and calcium-imaging techniques on rodents have emphasized the presence of notorious motor and arousal-related information in areas traditionally thought of as sensory (Vinck et al., 2015; Stringer et al., 2019). On the other hand, it may be argued from a conservative take on systems neurophysiology that primary visual areas are not particularly concerned with constructing face or affect percepts. Nonetheless, the classifier could have picked up and leveraged lower-order information in those areas to construct a statistical model about facial expression; similar to how artificial vision systems emulate cortical computations starting from nothing but raw pixels. That would certainly pose a methodological challenge to our approach (sensitivity in excess can be detrimental), which we showed was alleviated to some extent by the diligent use of control stimuli (neutral faces).

This result is of great interest, in light of the incipient works on emotion as seen through the MVPA prism. For instance some of the literature from table 1 also included anger and sadness-loaded stimuli in their experiments, reportedly with better results than ours (Ethofer et al., 2009; Said et al., 2010; Kotz et al., 2012). However, reasons exist to be skeptical of them upon closer inspection. The ROI-based, auditory study by Ethofer et al., for example, reported average classification accuracies (n=22) of 30% and >35% for sadness and anger respectively; among 5 emotions. Nonetheless, models were trained only pair-wisely: that is, contrasting target emotion against an “everything else” metaclass, which implies that performance was actually below the true chance level (50%). This one-vs-all scheme without nonparametric testing was repeated by Kotz et al., yet, here anger showed the poorest results. Other issues in the literature include comparing against a null model built from a scarce number of permutations, for instance, like in the study conducted by Said et al (2010).

Perhaps our results for these two basic emotions could have improved, had a more localized search been performed. Those affective-perceptual states might be genuinely underrepresented in the coarse fMRI data, or they may not be linearly separable, or the system dynamics may not be sufficiently stationary in the relevant dimensions. As argued during the Introduction, this study aimed at testing the limits of linear SVM as a data-driven anatomical mapping tool, at the expense of maximum decoding performance. In that sense, and joined by our modest sample size, being able to retrieve just *some* emotional states out of BOLD activity emanating from well-defined structures already cements the accomplishment of our goals.

The present work also suffers from limitations and future opportunity areas. Further analysis is required to characterize the intrinsic dimensionality of each cluster system. Similarly, system dynamics could be studied and modeled to provide further understanding of each successfully decoded state, as well as the encompassing attractor set. It would also be interesting to extend the task to other emotions, modalities and theoretical models of emotion; for instance to identify whether we have a sufficient characterization of happiness (as opposed to appetitive hedonic valence more generally, as posited by dimensional theories of emotion). A second strand of further studies could explore these findings using more direct causal interventions in the brain, so as to assess the relevance of the multivariate correlations we report.

## Acknowledgements

We are extremely grateful to Luis Concha, Eduardo Garza-Villarreal, Sarael Alcauter, Zeus Gracia-Tabuenca and Victor Olalde-Mathieu for their invaluable methodological guidance. To Eric Pasaye and UNAM’s MR unit personnel, to Victor Olalde-Mathieu, Ana Y. Martínez and Lluviana Rodríguez-Vidal for their help with data acquisition. We would also like to thank Leopoldo González-Santos, Aliza Brzezinski, Diego Ramírez and Enrique Chiu-Han for their general advice and technical assistance, and J.G. Norris for editing the manuscript.

## Supplementary material 1

### Behavioral results

Prior to entering the MRI machine, 15 of the 16 participants were asked to answer a brief randomized stimuli categorization task using the same faces they would later experiment inside the scanner. Faces could be assigned to one of four classes with a computer mouse: angry, happy, neutral or sad. A Pearson’s *χ*^2^ test for association strength between intended emotion and subjective interpretation assigned a probability of 6.9 *⋅* 10*^−^*^283^ to the possibility that participants were categorizing stimuli at random. A similar test was performed separating by stimulus as opposed to emotion class; nonetheless, the p-value remained very low at 3.2 *⋅* 10*^−^*^225^. Both results are shown in Figure 1. Despite variability recognizing among different basic emotions, our success rates turn out to be similar to those reported in independent validations of other datasets (Tottenham et al., 2009; Conley et al., 2018). Similarly, per-participant *χ*^2^ tests (with Bonferroni correction for FWE) revealed that even the worst-performing participant had a probability of less than 5 *⋅* 10*^−^* ^8^ of being involved in guesswork.

**Figure 1:**
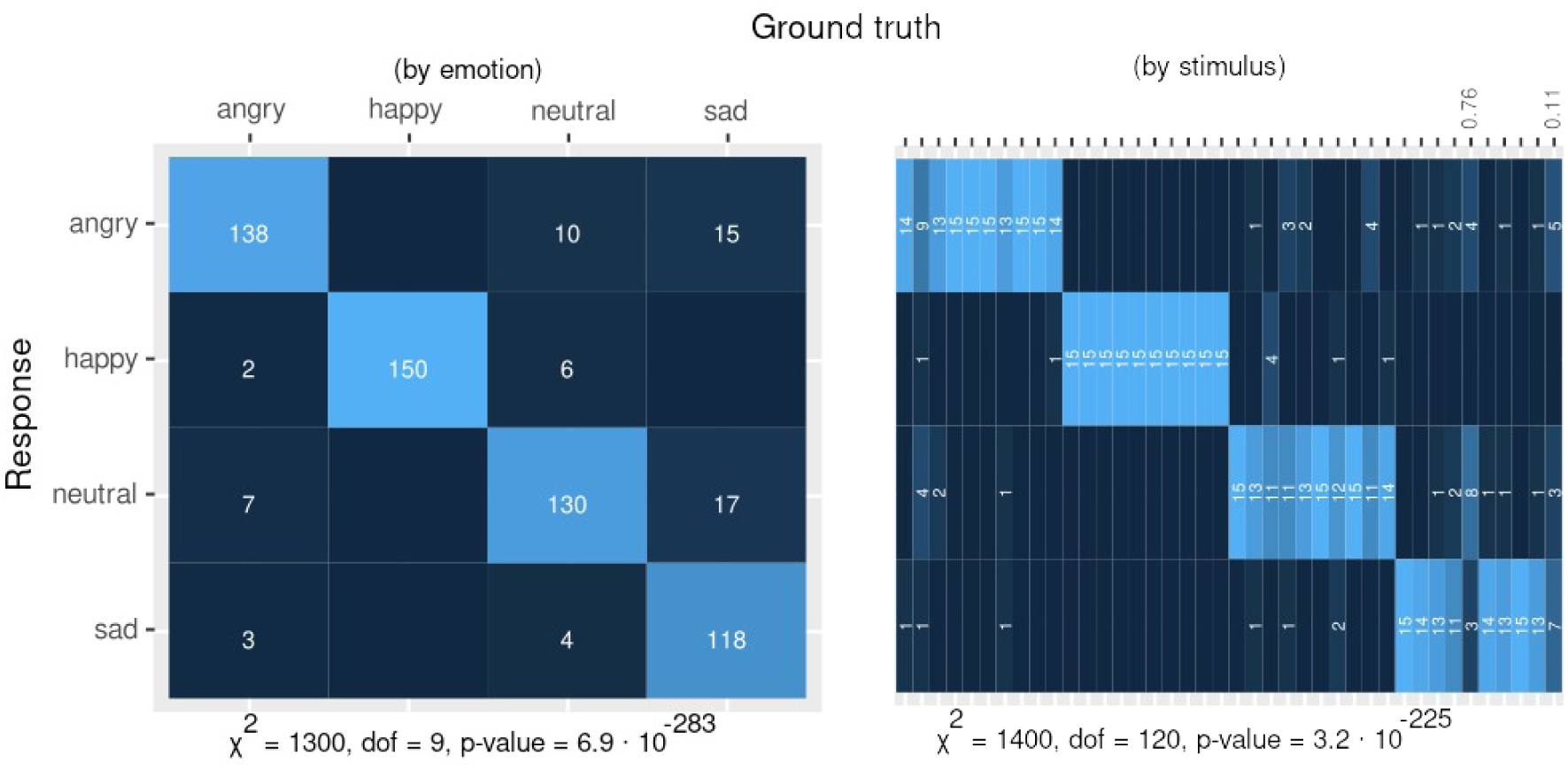
Confusion matrices with the group joint frequency of responses obtained during preparatory picture validation. For datasets true to their purpose, a strong diagonal should be observed, indicating agreement between subjective perception and preset categories. This is quantified with Pearson’s _χ_^2^ tests, whose results are displayed below each contingency table. **Top**: when grouping stimuli by emotion. **Bottom**: grouping only by picture, to detail the fine-grained structure of errors, Holm-corrected p-values are shown for the only two stimuli with p>0.05, according to one-tail binomial tests under the null hypothesis that the correct category is only assigned to 1/4th of all Bernoulli trials, presupposed to be statistically independent.

With regard to instantaneous responses during the task, we ran binomial tests to quantify success probability detecting face gender or image change, assuming statistical independence and a chance level of 50%. Figure 2 shows the aggregate of hits through time. Errors, in red, are comparatively low. A probability of 1.95 *⋅* 10*^−^*^60^ (Holm-corrected) of finding such hits/misses ratio by chance was found for the worst participant, and the probability for the worst block type (including pseudo-faces and dim-stimulation) is even smaller.

**Figure 2:**
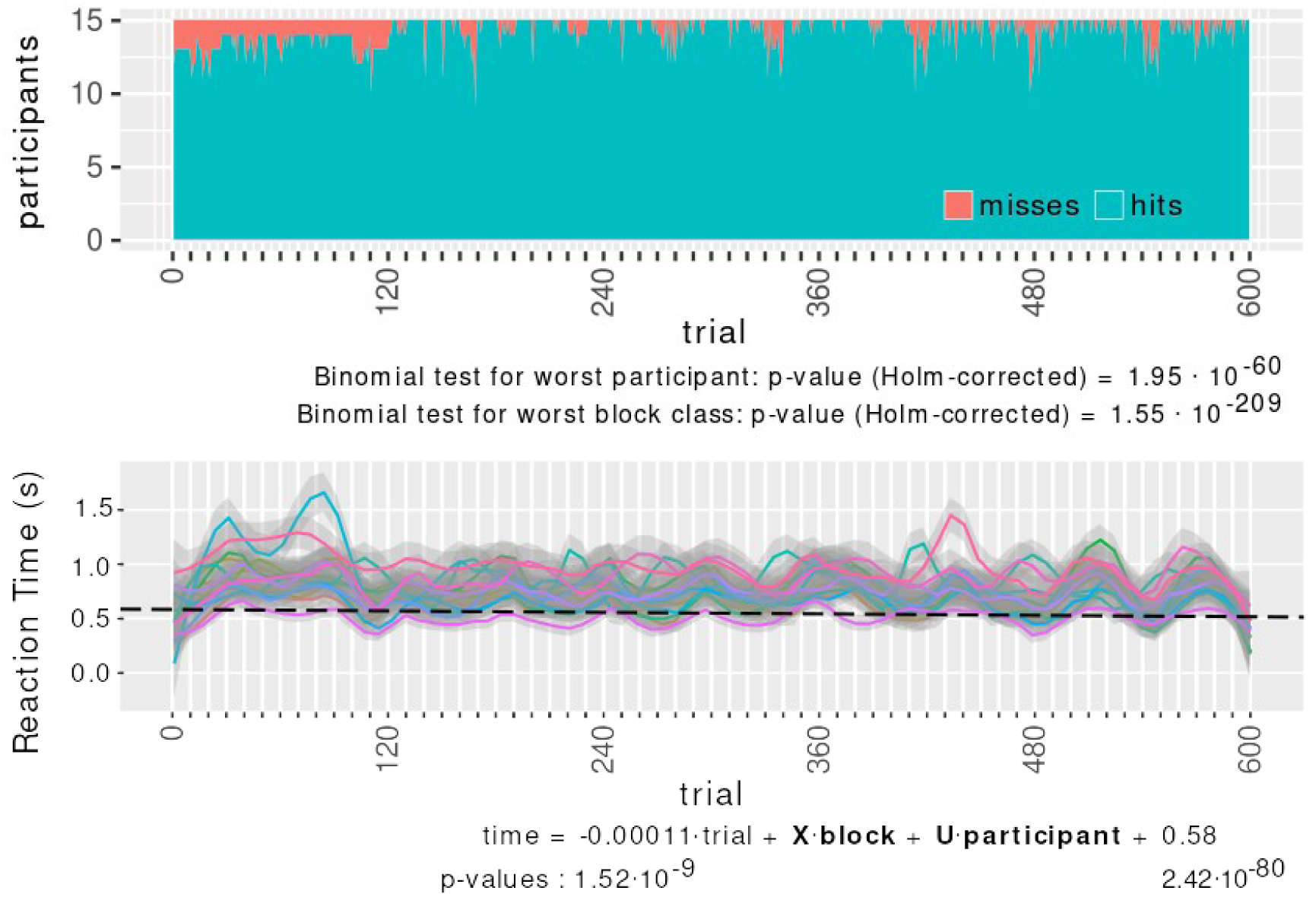
Instantaneous performance during the task. Each notch in the horizontal axis represents 10 consecutive trials; i.e., a 30 s block. Each sequence is delimited by the 120-trial-long marks. **Top**: ratio between hits and misses as a function of time. **Bottom**: per-subject reaction time, and a linear fit according to a GLMM. Each solid curve stands for the LOESS polynomial fit of a different participant and its confidence interval at 95%. The dotted line shows the almost-null linear trend from the GLMM. The actual GLMM is shown underneath, with explicit values for important parameters together with their respective p-values.

Participant’s reaction times (RTs) were analyzed as well, as a measure of attention to the task. Each curve in Figure 2 corresponds to the RTs of some subject. The superimposed dotted black line projects the relevant part of a general linear mixed-effects model (GLMM). GLMM is a generalization of GLM regression which uses two — as opposed to one — design matrices to account for random effects. This is specially suitable to hierarchical factorial designs; since the variance of measurements at some time *t* could come from intrinsic differences among participants, whose personal variance is captured by the random effects. The model was fitted using the participant factor as a random effect, and block lapse and block class as fixed effects. The aim is to find the effect of time upon RTs, because big changes would signify disengagement from the task. On the contrary, we observed a negligible, downward slope (0.11 ms faster RTs every 30 s block).

Moreover, a post-hoc Tukey test for a one-way ANOVA of RTs was inspected, using block types as factor levels. The only emotion to elicit considerably different reaction times was anger (angry vs neutral p=.014, angry vs happy p=.03). No significant difference was found between reacting to pseudo-faces vs to fixation crosses. However, we measured extremely large differences between reacting to any type of visuofacial stimulus and any type of non-visuofacial stimuli, for which not only stimulus complexity is lower, but task complexity is also lower (telling gender vs noticing any change at all).

All these lines of behavioral evidence converge towards the conclusion that participants understood the task and that stimuli were correctly observed in general. Accordingly, no participant or block type was discarded for analysis of the fMRI data after this screening.

## References

1. Amaral DG, Price J. (1984). Amygdalo-cortical projections in the monkey (macaca fascicularis). Journal of Comparative Neurology 230(4):465–96.

2. Amrhein, V., & Greenland, S. (2018). Remove, rather than redefine, statistical significance. Nature human behaviour, 2(1), 4–4.

3. Avena-Koenigsberger, A., Misic, B., & Sporns, O. (2018). Communication dynamics in complex brain networks. Nature reviews neuroscience, 19(1), 17–33.

4. Baucom LB, Wedell DH, Wang J, Blitzer DN, Shinkareva SV. (2012). Decoding the neural representation of affective states. Neuroimage 59(1):718–27.

5. Björnsdotter M, Rylander K, Wessberg J. (2011). A Monte Carlo method for locally multivariate brain mapping. Neuroimage 56(2):508–16.

6. Boser BE, Guyon IM, Vapnik VN (1992). A training algorithm for optimal margin classifiers. In: *Proceedings of the fifth annual workshop on computational learning theory* pp. 144–52.

7. Carlson TA, Schrater P, He S. (2003). Patterns of activity in the categorical representations of objects. Journal of cognitive neuroscience 15(5):704–17.

8. Celeghin A, Diano M, Bagnis A, Viola M, Tamietto M. (2017). Basic emotions in human neuroscience: Neuroimaging and beyond. Frontiers in Psychology 8:1432.

9. Chang LJ, Gianaros PJ, Manuck SB, Krishnan A, Wager TD. (2015). A sensitive and specific neural signature for picture-induced negative affect. PLoS biology 13(6):e1002180.

10. Chialvo, D. R. (2010). Emergent complex neural dynamics. Nature physics, 6(10), 744–750.

11. Chikazoe J, Lee DH, Kriegeskorte N, Anderson AK. (2014). Population coding of affect across stimuli, modalities and individuals. Nature neuroscience 17(8):1114.

12. Conley MI, Dellarco DV, Rubien-Thomas E, Cohen AO, Cervera A, Tottenham N, Casey BJ. (2018). The racially diverse affective expression (RADIATE) face stimulus set. Psychiatry research 270:1059–67.

13. Cox DD, Savoy RL. (2003). Functional magnetic resonance imaging (fMRI)“brain reading”: Detecting and classifying distributed patterns of fMRI activity in human visual cortex. Neuroimage 19(2):261–70.

14. Craddock, R. C., Holtzheimer III, P. E., Hu, X. P., & Mayberg, H. S. (2009). Disease state prediction from resting state functional connectivity. Magnetic Resonance in Medicine: An Official Journal of the International Society for Magnetic Resonance in Medicine, 62(6), 1619–1628.

15. Damaraju E, Huang Y-M, Barrett LF, Pessoa L. (2009). Affective learning enhances activity and functional connectivity in early visual cortex. Neuropsychologia 47(12):2480–7.

16. De Martino F, Valente G, Staeren N, Ashburner J, Goebel R, Formisano E. (2008). Combining multivariate voxel selection and support vector machines for mapping and classification of fMRI spatial patterns. Neuroimage 43(1):44–58.

17. Desikan, R. S., Ségonne, F., Fischl, B., Quinn, B. T., Dickerson, B. C., Blacker, D., … & Killiany, R.J. (2006). An automated labeling system for subdividing the human cerebral cortex on MRI scans into gyral based regions of interest. Neuroimage, 31(3), 968–980.

18. Ekman P. (1976). Pictures of facial affect. Consulting Psychologists Press.

19. Ethofer T, Van De Ville D, Scherer K, Vuilleumier P. (2009). Decoding of emotional information in voice-sensitive cortices. Current Biology 19(12):1028–33.

20. Fonov VS, Evans AC, McKinstry RC, Almli C, Collins D. (2009). Unbiased nonlinear average age-appropriate brain templates from birth to adulthood. NeuroImage (47):S102.

21. Fonov, V., Evans, A. C., Botteron, K., Almli, C. R., McKinstry, R. C., Collins, D. L., & Brain Development Cooperative Group. (2011). Unbiased average age-appropriate atlases for pediatric studies. Neuroimage, 54(1), 313–327.

22. Friston KJ, Holmes AP, Worsley KJ, Poline J-P, Frith CD, Frackowiak RS. (1994). Statistical parametric maps in functional imaging: A general linear approach. Human brain mapping 2(4):189–210.

23. Friston KJ, Frith CD, Frackowiak R, Turner R, others. (1995). Characterizing dynamic brain responses with fMRI: A multivariate approach. Neuroimage 2(2):166–72.

24. Gaonkar, B., & Davatzikos, C. (2013). Analytic estimation of statistical significance maps for support vector machine based multi-variate image analysis and classification. Neuroimage, 78, 270–283.

25. Guillory SA, Bujarski KA. (2014). Exploring emotions using invasive methods: Review of 60 years of human intracranial electrophysiology. Social cognitive and affective neuroscience 9(12):1880–9.

26. Haist, F., & Anzures, G. (2017). Functional development of the brain’s face-processing system. Wiley Interdisciplinary Reviews: Cognitive Science, 8(1-2), e1423.

27. Hamann S. (2012). Mapping discrete and dimensional emotions onto the brain: Controversies and consensus. Trends in cognitive sciences 16(9):458–66.

28. Hanke M, Halchenko YO, Sederberg PB, Hanson SJ, Haxby JV, Pollmann S. (2009). PyMVPA: A python toolbox for multivariate pattern analysis of fMRI data. Neuroinformatics 7(1):37–53.

29. Haxby, J. V., Hoffman, E. A., & Gobbini, M. I. (2000). The distributed human neural system for face perception. Trends in cognitive sciences, 4(6), 223–233.

30. Haynes J-D, Rees G. (2005). Predicting the orientation of invisible stimuli from activity in human primary visual cortex. Nature neuroscience 8(5):686–91.

31. Hoel EP, Albantakis L, Tononi G. (2013). Quantifying causal emergence shows that macro can beat micro. Proceedings of the National Academy of Sciences 110(49):19790–5.

32. Huettel SA, Song AW, McCarthy G. (2009). Functional magnetic resonance imaging. 2nd ed. Vol. 1. Sunderland, MA: Sinauer Associates.

33. Isenberg, N., Silbersweig, D., Engelien, A., Emmerich, S., Malavade, K., Beattie, B., … & Stern, E. (1999). Linguistic threat activates the human amygdala. Proceedings of the National Academy of Sciences, 96(18), 10456–10459.

34. Jenkinson M, Smith S. (2001). A global optimisation method for robust affine registration of brain images. Medical image analysis 5(2):143–56.

35. Jenkinson M, Bannister P, Brady M, Smith S. (2002). Improved optimization for the robust and accurate linear registration and motion correction of brain images. Neuroimage 17(2):825–41.

36. Jenkinson M, Beckmann CF, Behrens TE, Woolrich MW, Smith SM. (2012). FSL. Neuroimage 62(2):782–90.

37. Joliot, M., Jobard, G., Naveau, M., Delcroix, N., Petit, L., Zago, L., … & Tzourio-Mazoyer, N. (2015). AICHA: An atlas of intrinsic connectivity of homotopic areas. Journal of neuroscience methods, 254, 46–59.

38. Jolly, E., & Chang, L. J. (2021). Multivariate spatial feature selection in fMRI. Social Cognitive and Affective Neuroscience, 16(8), 795–806.

39. Kamitani Y, Tong F. (2005). Decoding the visual and subjective contents of the human brain. Nature neuroscience 8(5):679–85.

40. Kassam KS, Markey AR, Cherkassky VL, Loewenstein G, Just MA. (2013). Identifying emotions on the basis of neural activation. PloS one 8(6):e66032.

41. Kehoe, E. G., Toomey, J. M., Balsters, J. H., & Bokde, A. L. (2013). Healthy aging is associated with increased neural processing of positive valence but attenuated processing of emotional arousal: an fMRI study. Neurobiology of aging, 34(3), 809–821.

42. Kherif F, Poline J-B, Flandin G, Benali H, Simon O, Dehaene S, Worsley KJ. (2002). Multivariate model specification for fMRI data. Neuroimage 16(4):1068–83.

43. Kjems U, Hansen LK, Anderson J, Frutiger S, Muley S, Sidtis J, Rottenberg D, Strother SC. (2002). The quantitative evaluation of functional neuroimaging experiments: Mutual information learning curves. NeuroImage 15(4):772–86.

44. Kotz SA, Kalberlah C, Bahlmann J, Friederici AD, Haynes J-D. (2012). Predicting vocal emotion expressions from the human brain. Human Brain Mapping 34(8):1971–81.

45. Kragel PA, LaBar KS. (2014). Advancing emotion theory with multivariate pattern classification. Emotion Review 6(2):160–74.

46. Kragel PA, LaBar KS. (2015). Multivariate neural biomarkers of emotional states are categorically distinct. Social cognitive and affective neuroscience 10(11):1437–48.

47. Kragel PA, LaBar KS. (2016). Decoding the nature of emotion in the brain. Trends in cognitive sciences 20(6):444– 55.

48. Kriegeskorte N, Goebel R, Bandettini P. (2006). Information-based functional brain mapping. Proceedings of the National Academy of Sciences 103(10):3863–8.

49. Kriegeskorte N, Simmons WK, Bellgowan PS, Baker CI. (2009). Circular analysis in systems neuroscience: The dangers of double dipping. Nature neuroscience 12(5):535.

50. LaConte S, Anderson J, Muley S, Ashe J, Frutiger S, Rehm K, Hansen LK, Yacoub E, Hu X, Rottenberg D, Strother S. (2003). The evaluation of preprocessing choices in single-subject bold fMRI using npairs performance metrics. NeuroImage 18(1):10–27.

51. LaConte S, Strother S, Cherkassky V, Anderson J, Hu X. (2005). Support vector machines for temporal classification of block design fMRI data. NeuroImage 26(2):317–29.

52. Lewis-Peacock JA, Norman KA. (2013). Multi-voxel pattern analysis of fMRI data. The cognitive neurosciences 911–20.

53. Lindquist KA, Wager TD, Kober H, Bliss-Moreau E, Barrett LF. (2012). The brain basis of emotion: A meta-analytic review. Behavioral and brain sciences 35(3):121–43.

54. Manjón JV, Coupé P. VolBrain: An online MRI brain volumetry system. (2016). Frontiers in neuroinformatics 10:30.

55. Mahmoudi A, Takerkart S, Regragui F, Boussaoud D, Brovelli A. (2012). Multivoxel pattern analysis for FMRI data: A review. Computational and mathematical methods in medicine.

56. McIntosh A, Bookstein F, Haxby JV, Grady C. (1996). Spatial pattern analysis of functional brain images using partial least squares. Neuroimage 3(3):143–57.

57. McKeown MJ, Makeig S, Brown GG, Jung T-P, Kindermann SS, Bell AJ, Sejnowski TJ. (1998). Analysis of fMRI data by blind separation into independent spatial components. Human brain mapping 6(3):160–88.

58. Mørch N, Hansen LK, Strother SC, Svarer C, Rottenberg DA, Lautrup B, Savoy R, Paulson OB. Nonlinear versus linear models in functional neuroimaging: Learning curves and generalization crossover. (1997). In: *Biennial international conference on information processing in medical imaging*. Springer pp. 259–70.

59. Mourão-Miranda J, Bokde AL, Born C, Hampel H, Stetter M. (2005). Classifying brain states and determining the discriminating activation patterns: Support vector machine on functional MRI data. NeuroImage 28(4):980–95.

60. Mwangi B, Tian TS, Soares JC. (2014). A review of feature reduction techniques in neuroimaging. Neuroinformatics 12(2):229–44.

61. Palo, H. K., Sahoo, S., & Subudhi, A. K. (2021). Dimensionality reduction techniques: Principles, benefits, and limitations. Data Analytics in Bioinformatics: A Machine Learning Perspective, 77–107.

62. Peelen MV, Atkinson AP, Vuilleumier P. (2010). Supramodal representations of perceived emotions in the human brain. Journal of Neuroscience 30(30):10127–34.

63. Peirce JW. (2007). PsychoPy—psychophysics software in python. Journal of neuroscience methods 162(1-2):8–13.

64. Pessoa L, Padmala S. (2007). Decoding near-threshold perception of fear from distributed single-trial brain activation. Cerebral cortex 17(3):691–701.

65. Polyn, S. M., Natu, V. S., Cohen, J. D., & Norman, K. A. (2005). Category-specific cortical activity precedes retrieval during memory search. Science, 310(5756), 1963–1966.

66. Prinz, J. (2006). Is the mind really modular. Contemporary debates in cognitive science, 14, 22–36.

67. Raizada RD, Tsao F-M, Liu H-M, Holloway ID, Ansari D, Kuhl PK. (2010). Linking brain-wide multivoxel activation patterns to behaviour: Examples from language and math. NeuroImage 51(1):462–71.

68. Roiser J, Linden D, Gorno-Tempinin M, Moran R, Dickerson B, Grafton S. (2016). Minimum statistical standards for submissions to neuroimage: Clinical. NeuroImage: Clinical 12:1045.

69. Rolls ET, Grabenhorst F, Franco L. (2009). Prediction of subjective affective state from brain activations. Journal of Neurophysiology 101(3):1294–308.

70. Saarimäki H, Gotsopoulos A, Jääskeläinen IP, Lampinen J, Vuilleumier P, Hari R, Sams M, Nummenmaa L. (2015). Discrete neural signatures of basic emotions. Cerebral cortex 26(6):2563–73.

71. Said CP, Moore CD, Engell AD, Todorov A, Haxby JV. (2010). Distributed representations of dynamic facial expressions in the superior temporal sulcus. Journal of vision 10(5):11–1.

72. Schmah T, Yourganov G, Zemel RS, Hinton GE, Small SL, Strother SC. (2010). Comparing classification methods for longitudinal fMRI studies. Neural computation 22(11):2729–62.

73. Schmahmann JD, Sherman JC. (1997). Cerebellar cognitive affective syndrome. International review of neurobiology 41:433–40.

74. Sejnowski TJ. (2020). The unreasonable effectiveness of deep learning in artificial intelligence. Proceedings of the National Academy of Sciences. 117(48), 30033–30038.

75. Shinkareva SV, Wang J, Kim J, Facciani MJ, Baucom LB, Wedell DH. (2014). Representations of modality-specific affective processing for visual and auditory stimuli derived from functional magnetic resonance imaging data. Human brain mapping 35(7):3558–68.

76. Sitaram R, Lee S, Ruiz S, Rana M, Veit R, Birbaumer N. (2011). Real-time support vector classification and feedback of multiple emotional brain states. Neuroimage 56(2):753–65.

77. Skerry AE, Saxe R. (2015). Neural representations of emotion are organized around abstract event features. Current biology 25(15):1945–54.

78. Smith SM, Nichols TE. (2009). Threshold-free cluster enhancement: Addressing problems of smoothing, threshold dependence and localisation in cluster inference. Neuroimage 44(1):83–98.

79. Stringer C, Pachitariu M, Steinmetz N, Reddy CB, Carandini M, Harris KD. (2019). Spontaneous behaviors drive multidimensional, brainwide activity. Science 364(6437):eaav7893.

80. Tottenham N, Tanaka JW, Leon AC, McCarry T, Nurse M, Hare TA, Marcus DJ, Westerlund A, Casey BJ, Nelson C. (2009). The NimStim set of facial expressions: Judgments from untrained research participants. Psychiatry research 168(3):242–9.

81. Tzourio-Mazoyer, N., Landeau, B., Papathanassiou, D., Crivello, F., Etard, O., Delcroix, N., … & Joliot, M. (2002). Automated anatomical labeling of activations in SPM using a macroscopic anatomical parcellation of the MNI MRI single-subject brain. Neuroimage, 15(1), 273–289.

82. Vapnik V, Chervonenkis A. (1974). Theory of pattern recognition. Moscow. Nauka.

83. Vinck M, Batista-Brito R, Knoblich U, Cardin JA. (2015). Arousal and locomotion make distinct contributions to cortical activity patterns and visual encoding. Neuron 86(3):740–54.

84. Vytal K, Hamann S. (2010). Neuroimaging support for discrete neural correlates of basic emotions: A voxel-based meta-analysis. Journal of cognitive neuroscience 22(12):2864–85.

85. Wang, Z., Childress, A. R., Wang, J., & Detre, J. A. (2007). Support vector machine learning-based fMRI data group analysis. NeuroImage, 36(4), 1139–1151.

86. Wasserstein, R. L., & Lazar, N. A. (2016). The ASA statement on p-values: context, process, and purpose. The American Statistician, 70(2), 129–133.

87. Wasserstein, R. L., Schirm, A. L., & Lazar, N. A. (2019). Moving to a world beyond “p< 0.05”. The American Statistician, 73(sup1), 1–19.

88. Wegrzyn M, Riehle M, Labudda K, Woermann F, Baumgartner F, Pollmann S, Bien C, Kissler J. (2015). Investigating the brain basis of facial expression perception using multi-voxel pattern analysis. Cortex 69:131–40.

89. Winkler AM, Ridgway GR, Webster MA, Smith SM, Nichols TE. (2014). Permutation inference for the general linear model. Neuroimage 92:381–97.

90. Woolrich MW, Ripley BD, Brady M, Smith SM. (2001). Temporal autocorrelation in univariate linear modeling of FMRI data. Neuroimage 14(6):1370–86.

91. Zhang J, Zhang G, Li X, Wang P, Wang B, Liu B. (2020). Decoding sound categories based on whole-brain functional connectivity patterns. Brain imaging and behavior 14(1):100–9.

